# Overlapping but asymmetrical relationships between schizophrenia and autism revealed by brain connectivity

**DOI:** 10.1101/403212

**Authors:** Yujiro Yoshihara, Giuseppe Lisi, Noriaki Yahata, Junya Fujino, Yukiko Matsumoto, Jun Miyata, Genichi Sugihara, Shin-ichi Urayama, Manabu Kubota, Masahiro Yamashita, Ryuichiro Hashimoto, Naho Ichikawa, Weipke Cahn, Neeltje E. M. van Haren, Susumu Mori, Yasumasa Okamoto, Kiyoto Kasai, Nobumasa Kato, Hiroshi Imamizu, René S. Kahn, Akira Sawa, Mitsuo Kawato, Toshiya Murai, Jun Morimoto, Hidehiko Takahashi

**Affiliations:** Department of Psychiatry, Graduate School of Medicine, Kyoto University, Kyoto 606-8507, Japan; Department of Brain Robot Interface, ATR (Advanced Telecommunications Research Institute International) Brain Information Communication Research Laboratory Group, Kyoto 619-0288, Japan; Department of Decoded Neurofeedback, ATR Brain Information Communication Research Laboratory Group, Kyoto 619–0288, Japan; Department of Youth Mental Health, Graduate School of Medicine, The University of Tokyo, Tokyo 113–0033, Japan; Department of Molecular Imaging and Theranostics, National Institute of Radiological Sciences, National Institutes for Quantum and Radiological Science and Technology, Chiba 263–8555, Japan; Medical Institute of Developmental Disabilities Research, Showa University Karasuyama Hospital, Tokyo 157-8577, Japan; Human Brain Research Center, Graduate School of Medicine, Kyoto University, Kyoto 606- 8507, Japan; Department of Functional Brain Imaging Research, National Institute of Radiological Sciences, National Institutes for Quantum and Radiological Science and Technology, Chiba 263-8555, Japan; Department of Cognitive Neuroscience, ATR Brain Information Communication Research Laboratory Group, Kyoto 619-0288, Japan; Department of Language Sciences, Tokyo Metropolitan University, Tokyo 192-0397, Japan; Department of Psychiatry and Neurosciences, Graduate School of Biomedical Sciences, Hiroshima University, Hiroshima 734-8553, Japan; Department of Psychiatry, Brain Centre Rudolf Magnus, University Medical Center Utrecht, Heidelberglaan 100, 3584 CX Utrecht, The Netherlands; Department of Radiology, Johns Hopkins University School of Medicine, Baltimore, Maryland 21205, USA; Department of Neuropsychiatry, Graduate School of Medicine, The University of Tokyo, Tokyo 113-8655, Japan; Department of Psychology, Graduate School of Humanities and Sociology, The University of Tokyo, Tokyo 113-0033, Japan; Department of Psychiatry, Johns Hopkins University School of Medicine, Baltimore, Maryland 21205, USA

## Abstract

Although the relationship between schizophrenia spectrum disorder (SSD) and autism spectrum disorder (ASD) has long been debated, it has not yet been fully elucidated. To address this issue, we took advantage of dual (ASD and SSD) classifiers that discriminate patients from their controls based on resting state brain functional connectivity. An SSD classifier using sophisticated machine-learning algorithms that automatically selected SSD-specific functional connections was applied to Japanese datasets including adult patients with SSD in a chronic stage. We demonstrated good performance of the SSD classification for independent validation cohorts. The generalizability was tested by USA and European cohorts in a chronic stage, and one USA cohort including first episode schizophrenia. The specificity was tested by two adult Japanese cohorts of ASD and major depressive disorder, and one European cohort of attention-deficit hyperactivity disorder. The weighted linear summation of the classifier’s functional connections constituted the biological dimensions representing neural liability to the disorders. Our previously developed robust ASD classifier constituted the ASD dimension. Distributions of individuals with SSD, ASD and healthy controls were examined on the SSD and ASD biological dimensions. The SSD and ASD populations exhibited overlapping but asymmetrical patterns on the two biological dimensions. That is, the SSD population showed increased liability on the ASD dimension, but not vice versa. Furthermore, the two dimensions were correlated within the ASD population but not the SSD population. Using the two biological dimensions based on resting-state functional connectivity enabled us to quantify and visualize the relationships between SSD and ASD.

## Introduction

The relationship between schizophrenia and autism is a matter of historical and long-lasting debate. In 1911, Eugen Bleuler regarded autism as one of the fundamental symptoms in schizophrenia ^1^. No clear distinction between schizophrenia and autism had been described by the presentation of Diagnostic and Statistical Manual of Mental Disorders (DSM)-II in 1968. In the mid-60s to 70s, epidemiological studies concluded that these two conditions were distinct and unrelated. However, recent biological studies showed overlapping relationships and commonalities between the two disorders ^2, 3^. Genetic studies demonstrated common loci and pathways, suggesting that autism spectrum disorder (ASD) overlaps with schizophrenia spectrum disorder (SSD) ^4, 5^. Brain structural magnetic resonance imaging (MRI) and functional MRI studies also reported common abnormalities in gray matter volumes ^6^ and brain activations ^7, 8^. Nevertheless, the relationship between SSD and ASD remains controversial ^2^.

The fundamental problems behind this issue are that we lack a reliable biological identification for these disorders and that the diagnosis is based mostly on a symptomatological and categorical approach as represented by DSM. DSM criteria are mainly based on the patient’s behavioral signs and symptoms ^9^, although the symptoms in patients with SSD and ASD, respectively, are heterogeneous and vary erratically over time ^10, 11^. Hence, there is an explanatory gap between phenomenological entities and neurobiological underpinnings. To bridge this explanatory gap, researchers have begun to use a dimensional approach advocated by the National Institute of Mental Health Research Domain Criteria ^12^. The dimensional approach involves exploratory analysis with a vast amount of data ^13^, and therefore its constructs at the moment are distant from the level of “the actual clinical phenomena that bring patients to the clinic ^14, 15^”.

To solve the problems detailed above and to unravel the relationship between SSD and ASD, we propose a novel approach that reconciles categorical and dimensional approaches, that is, establishment of biological dimensions that are also compatible with DSM-based categorical diagnostic labels. We recruited individuals with ASD and SSD according to DSM. Next, we developed ASD and SSD classifiers using sophisticated machine-learning algorithms from brain functional connectivity (FC) measured by resting-state fMRI (rs-fMRI), based on the reports that ASD ^16-21^ and SSD ^22-29^ exhibited FC abnormalities in rs-fMRI. The classifiers for biological dimensions must be robust enough to have generalizability to independent cohorts with different ethnicities or MRI machine vendors. We have already developed the ASD classifier that has generalizability to perfectly independent validation cohorts ^30^, and here, we developed a similarly generalizable SSD classifier using the same machine-learning methods. Furthermore, we determined each biological dimension from the weighted linear summation of functional connections of SSD and ASD classifiers, and plotted individuals with ASD, SSD, and healthy controls (HCs) on the SSD-ASD dimensions. Finally, visualizing and quantifying each individual in a relative manner, we could verify the relationship between SSD and ASD populations.

## Methods

### Participants and MRI data acquisition

**Kyoto:** A total of 68 adult patients with SSD, including 64 patients with schizophrenia and 4 patients with schizoaffective disorder, and 102 HCs were recruited at the Department of Psychiatry, Kyoto University. We recruited two groups: Kyoto A and Kyoto B (Supplementary Methods and Table S1). At Kyoto A, we recruited 18 SSD, including 17 patients with schizophrenia and one patient with schizoaffective disorder, and 29 HCs. At Kyoto B, we recruited 50 SSD, including 47 patients with schizophrenia, 3 patients with schizoaffective disorder, and 73 HCs. All participants in the present study provided written informed consent that was approved by the Committee on Medical Ethics of Kyoto University. All patients were receiving antipsychotic medications. T1-structural and resting-state functional MR images at Kyoto A and B were scanned on 3T Siemens TimTrio and 3T Siemens Trio, respectively (Table S2).

### Preprocessing of MR images

MRI datasets (68 SSD and 102 HC) for training of the SSD/HC classifier in Kyoto were preprocessed, and calculation of a correlation matrix was performed using Statistical Parametric Mapping 8 (SPM8; Wellcome Trust Center for Neuroimaging, University College London, UK) software running on MATLAB (R2014a, Mathworks, USA) in the same manner as in our previous study ^30^ (Supplementary Methods).

### Selecting FCs as SSD classifier

To develop an SSD classifier from the correlation matrices, we adopted a cascade of *L*_1_-norm regularized sparse canonical correlation analysis (*L*_1_-SCCA) ^31^ and sparse logistic regression (SLR) ^32^ to select SSD-specific FCs while minimizing the effects of over-fitting and nuisance variables. The selection of SSD-specific FCs and classification performance evaluation were carried out through a sequential process of 9 x 9 nested feature-selection and leave-one-out cross-validation (LOOCV). The machine-learning algorithms automatically selected 10-20 FCs from about 10,000 FCs of whole brain rs-fMRI. The weighted linear summation (WLS) of the correlation values of the selected FCs predicted the categorical diagnostic label for each individual. The positive and negative values of WLS in each individual corresponded to a patient and a healthy control (HC), respectively. At the same time, a logistic regression function of WLS computes a probability for the patient. More concretely, the value of the continuous WLS provided a classification certainty. A large positive value indicates high certainty for SSD or ASD, a large negative value for high certainty for HC, and values near zero indicate uncertainty. Consequently, the WLS distributions based on functional brain connectivity probabilistically determined the neural liability to ASD and SSD as well as candidate genes for the disorders determined the genetic liability. Thus, we here named the WLS value “neural liability”. In the beginning of applying the machine-learning algorithm to patient and HC populations, we employed the binary value (patient or HC) of categorical diagnosis, and at the end of this process, we generated the continuous probabilistic degree of diagnostic certainty as an objective neural liability. Then, we utilized the neural liability as a biological dimension. In this way, we could integrate the categorical and dimensional approaches. (Figure S1 and Supplementary Methods).

The performance of the classifier was expressed in terms of area under the curve (AUC), accuracy, sensitivity, and specificity. The statistical significance of classification was assessed by permutation test ^33^.

### Generalizability of the Kyoto classifier

We tested the generalizability of the Kyoto classifier to three independent cohorts, COBRE of the Mind Research Network (Center for Biomedical Research Excellence, University of New Mexico, USA), UMCU-TOPFIT (The Outcome of Psychosis and Fitness Therapy, University Medical Centre Utrecht, The Netherlands), and a first episode schizophrenia cohort JHU-FES (Johns Hopkins University, USA) (Supplementary Methods and Table S3- S4). The patients with SSD of Kyoto, COBRE, and UMCU-TOPFIT were mainly in a chronic stage of the disease, while JHU-FES was in an early stage. The external datasets (COBRE, UMCU-TOPFIT, and JHU-FES) were preprocessed in the same manner as the Kyoto dataset.

**COBRE:** COBRE is the dataset publicly available at http://fcon_1000.projects.nitrc.org/indi/retro/cobre.html. A total of 46 patients with SSD, including 41 patients with schizophrenia and 5 patients with schizoaffective disorder, and 61 HCs were recruited. The ethnicity of most participants was Caucasian or Hispanic. 44 patients with SSD were receiving antipsychotic medications, one SSD was not receiving antipsychotics and one SSD had no information about the use of antipsychotics. The MR images of COBRE were scanned on 3T Siemens TimTrio.

**UMCU-TOPFIT:** A total of 47 patients with SSD, including 35 patients with schizophrenia and 12 patients with schizoaffective disorder, and 43 HCs were recruited. About four-fifths of the participants were born in the Netherlands. The patients in the UMCU-TOPFIT study were recruited at four different locations in The Netherlands. 44 patients with SSD were receiving antipsychotic medications and 3 patients with SSD were not. The MR images of UMCU- TOPFIT were scanned on 3T Philips Achieva.

**JHU-FES:** A total of 30 patients with FES, including 21 patients with schizophrenia, 7 with schizoaffective disorder, one with schizophreniform and one with psychotic disorder not otherwise specified, and 71 HCs were recruited at Johns Hopkins University hospital and incorporated into the present analysis. The ethnicity of many participants was African-American or Caucasian. 21 patients with FES were receiving antipsychotic medication, 3 patients with FES were not taking antipsychotics, and 6 patients with FES provided no information about any current use of antipsychotics. The MR images of JHU-FES were scanned on 3T Philips Achieva.

### Specificity of the Kyoto classifier

We tested the specificity of the Kyoto classifier, applying the classifier to two additional Japanese cohorts of ASD and major depressive disorder (MDD), respectively, and one European cohort of attention-deficit hyperactivity disorder (ADHD) (Supplementary Methods). The datasets of ASD, ADHD, and MDD were scanned on 3T MRI system. Details of their demographic information and MRI parameters are shown in the referred study ^30^. The other disorder’s datasets (ASD, ADHD, and MDD) were preprocessed in the same manner as the Kyoto dataset. The WLS distributions between each disorder population (ASD, ADHD, and MDD) and the corresponding HCs were compared via the AUC and Kolmogorov-Smirnov test.

**ASD:** A total of 74 adults with ASD and 107 age, sex, handedness and IQ-matched typically developed individuals as HCs were examined. 18 adults with ASD were receiving antipsychotic medications. The participants were recruited at three different locations (the University of Tokyo Hospital, Showa University Karasuyama Hospital, and Advanced Telecommunications Research Institute International) in Japan.

**ADHD:** A total of 13 participants with ADHD and age-matched 13 HCs were examined. The dataset was acquired by the NeuroIMAGE project in the Netherlands (http://www.neuroimage.nl/).

**MDD**: A total of 104 patients with MDD and 143 age-matched HC were examined. The patients were recruited from a local clinic and the healthy controls from the community of the Hiroshima University.

### Relationships between SSD and ASD on the two biological dimensions

Japanese individuals with SSD, ASD, and HC were plotted on the SSD-ASD dimensional plane. The SSD and ASD dimensional scores are the WLS using the SSD and ASD classifiers, respectively. The ASD classifier was taken from our previous study ^30^. The WLS distributions between each disorder population (ASD and SSD) and the corresponding HCs were compared via the AUC and Kolmogorov-Smirnov test. Categorical ellipses of SSD, ASD and HC were calculated using multivariate Gaussian distribution. The principal axes of the ellipses were obtained by computing the eigenvectors of the multivariate Gaussian’s covariance. Projections from the center of the clusters showed differences among their respective means.

To understand the impact of individual FCs on the ASD-SSD relationship, we analyzed the contribution to the WLS of the other disorder’s population (e.g. SSD) for each FC selected by the classifier of one disorder (e.g. ASD). The contribution of each FC within a population is computed by averaging the FC weighted by the classifier’s weight. A large positive difference between the disorder and control contributions indicates that a specific FC contributes positively to the classification.

Furthermore, separately for each population we analyzed the correlation coefficients between the most relevant FCs (top 5 each; 25 correlations) selected by the ASD and SSD classifiers and the cumulative sum across correlation coefficients in order to find the general trend of correlation. This was done separately for the two populations of patients. Before computing the correlation coefficients, the FCs are weighted by the sign of the classifier’s weight, in order to obtain positive correlation coefficients for the FCs contributing positively to the WLS. For a given population (e.g. ASD), one of the dimensions (e.g. ASD) is computed by LOOCV, while the other dimension (e.g. SSD) is computed by a one-shot prediction using the classifier built with the alternative disorder. For this reason, the most relevant FCs are defined as the top five FCs with largest cumulative absolute weight across cross-validation folds for LOOCV, and as the top five FCs with the largest absolute weight for the one-shot classifier.

## Results

### Accurate SSD classifier for Kyoto discovery cohort

The 16 FCs incorporated in our final classifier were selected by the sparse logistic regression (SLR) using the whole Kyoto datasets. The identified FCs showed the robustness and stability of across the cross-validation procedure (Figure S2). The classifier differentiated SSD from HC populations with an accuracy of 76% and an AUC of 0.83 (permutation test, *P* = 0.006; see Table 1 and Figure S3). We calculated the WLS of each participant from the 16 FCs. The two WLS distributions of the SSD and HC populations were clearly separated by a threshold of WLS = 0 (Fig. 1a). We found that high classification accuracy was not only achieved for the entire datasets, but also for the two sites separately (the accuracies of Kyoto A and B were 74% and 77%, respectively) (Table 1 and Figure S4). When tested on the COBRE dataset, the Kyoto classifier achieved high performance, with an accuracy of 70% (AUC=0.75) (Table 1 and Fig. 1b). The probability of obtaining this high performance by chance is as small as *P* = 0.001 (permutation test, see Figure S3). For UMCU-TOPFIT (Fig. 1c), the classifier also achieved accuracy of 61% (AUC=0.66) (*P* = 0.031, permutation test), although this classification performance for UMCU-TOPFIT was lower than for COBRE. For JHU-FES (Fig. 1d), the AUC (0.42) was below the chance level (Table 1), and thus generalization was not observed.

**Table 1.** Performance of the SSD classifier for the Kyoto, COBRE, UMCU-TOPFIT, and JHU-FES datasets

| Dataset | AUC | Accuracy<br>(%) | Sensitivity<br>(%) | Specificity<br>(%) | Accuracy<br>Kyoto A | Accuracy<br>Kyoto B |
| --- | --- | --- | --- | --- | --- | --- |
| Kyoto (N = 170) | 0.83 | 76 | 72 | 79 | 74 | 77 |
| COBRE (N = 107) | 0.75 | 70 | 65 | 74 |  |  |
| UMCU-TOPFIT (N = 90) | 0.66 | 61 | 64 | 58 |  |  |
| JHU-FES (N = 101) | 0.42 | 45 | 40 | 47 |  |  |

**Fig. 1.**
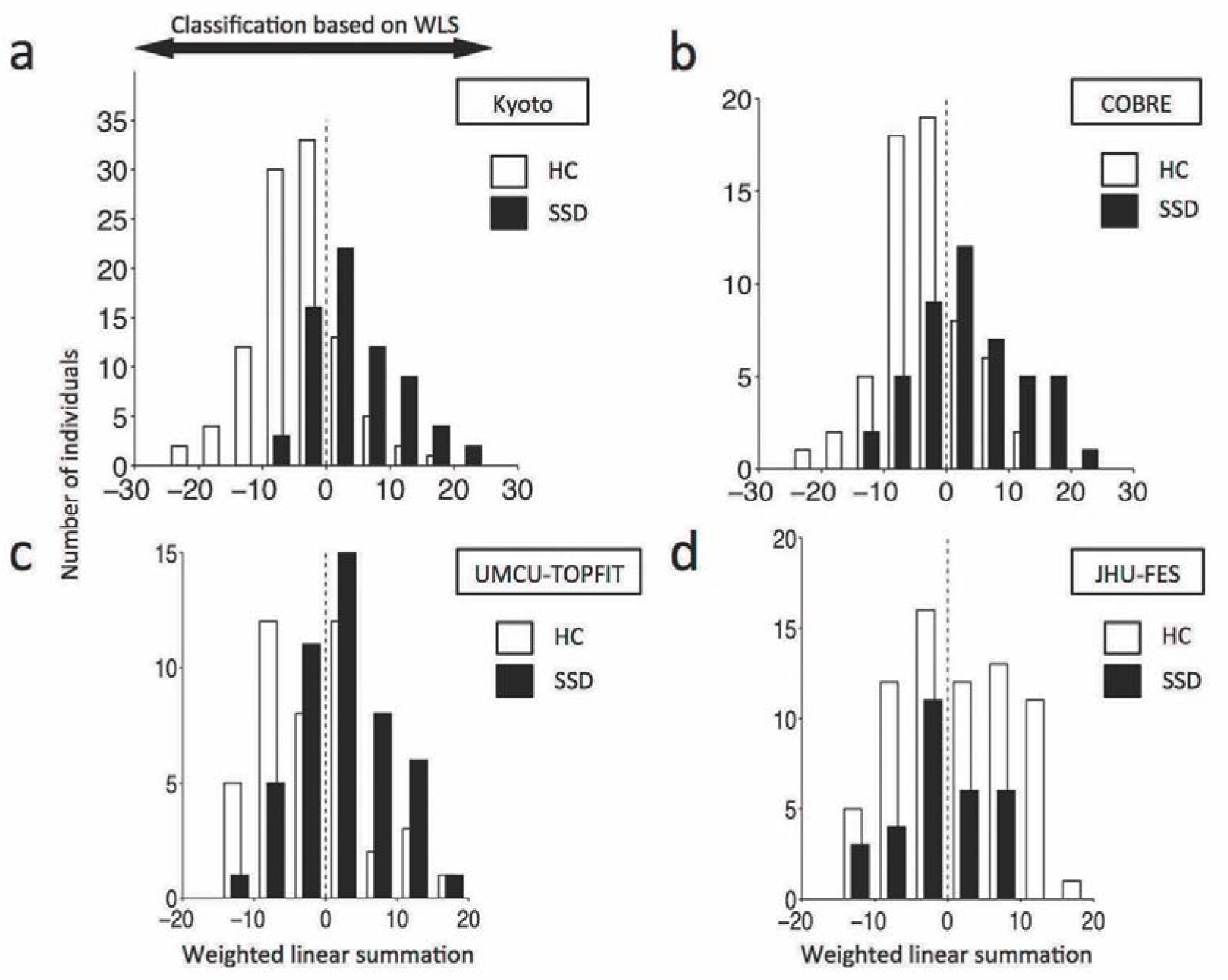
Distribution of weighted linear summation (WLS) of the SSD classifier. (**a**) The number of HC (white) and SSD (black) individuals in the Kyoto (Japan) data included in a specific WLS interval of width 5 is shown as a histogram. (**b, c, d**) WLS for the COBRE (USA), UMCU-TOPFIT (The Netherlands), and JHU-FES (USA) datasets are shown in the same formats as in (**a**). For this classifier, the WLS (or linear discriminant function) of the correlation values of the identified FC predicted the diagnostic label of each individual. A participant with positive or negative WLS was classified as SSD or HC, respectively.

### Characteristics of 16 identified FCs in the SSD classifier

The 16 FCs as SSD classifier were distributed as inter-hemispheric (44%), left intra-hemispheric (25%), and right intra-hemispheric connections (31%) (Fig. 2a-2b, Supplementary Results, and Table S5). The 16 FCs as SSD classifier were different from the 16 FCs as ASD classifier that we previously developed ^30^ (Fig. 2c and Figure S5).

**Fig. 2.**
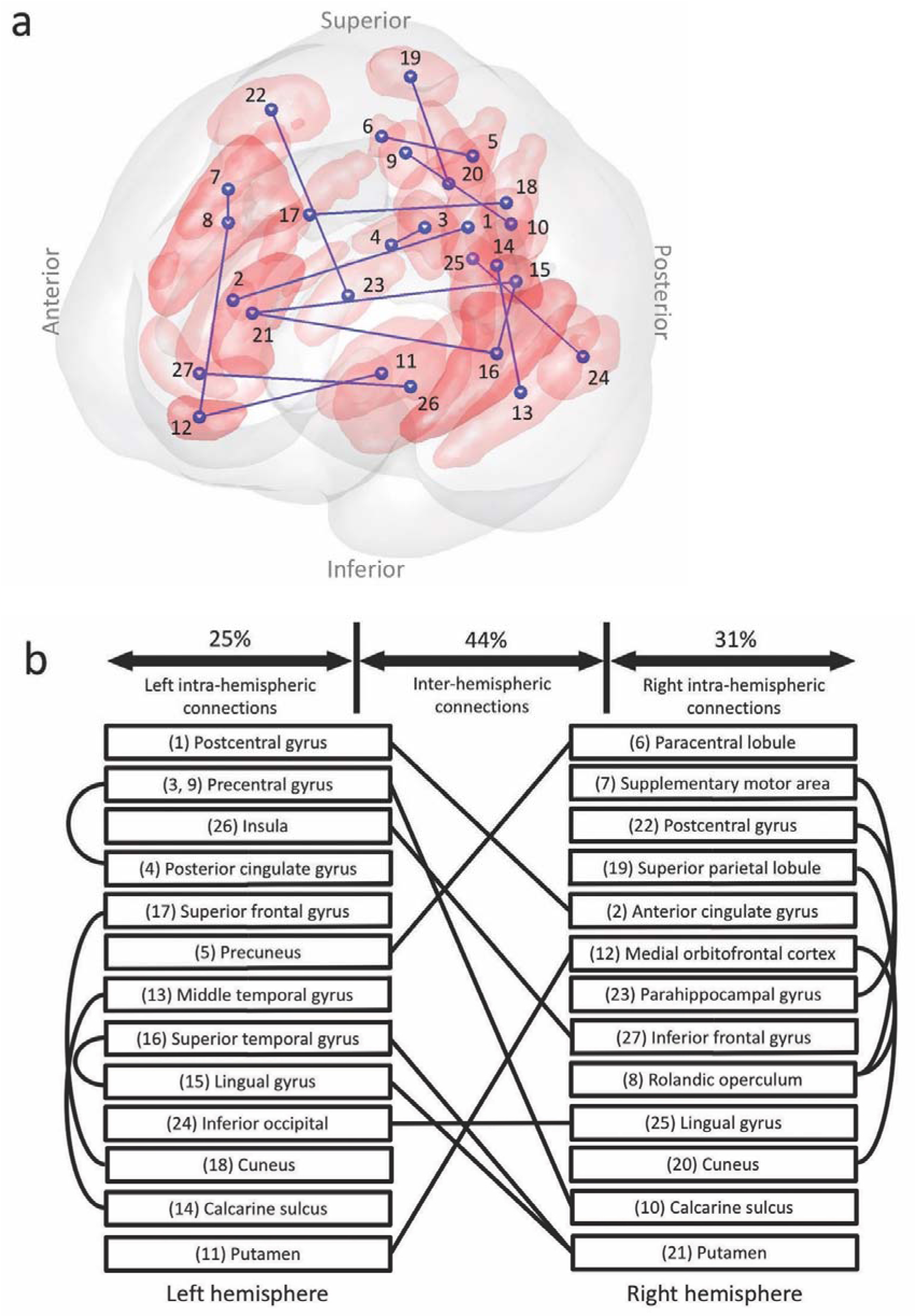
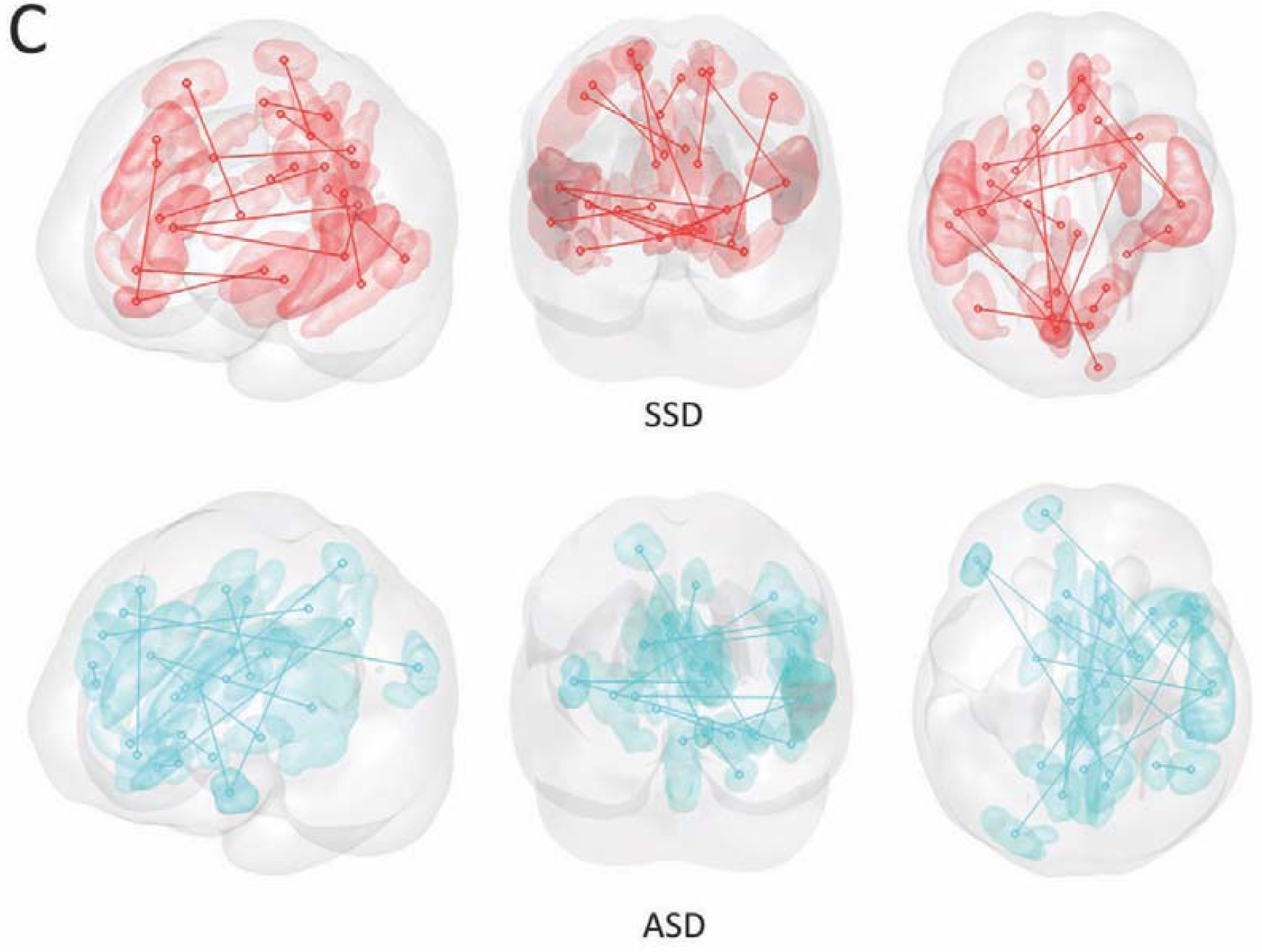
The 16 functional connectivities (FCs) of the SSD classifier. The 16 FCs viewed from anterior-left (**a**). (**b**) The 16 FCs (solid lines) and their terminal regions (names in boxes) are presented (details noted in Supplementary Results and Table S6). The left and right halves of the figure correspond to the left and right brain hemispheres, respectively. The FCs were classified into three hemispherical categories: left intra-hemispheric, right intra-hemispheric and inter-hemispheric. The terminal regions were defined by the anatomical automatic labelling (AAL). (**c**) The 16 FCs as SSD classifier (red lines and areas) were entirely different from the 16 FCs as ASD classifier (cyan lines and areas).

### Specificity of the classifier to SSD regarding other psychiatric disorders

Separation of WLS distribution was largest between SSD and HC (Fig. 3a) as already shown (Fig. 1a). In ASD, ADHD and MDD, the distribution was not distinguishable from HC (AUC = 0.50, Kolmogorov-Smirnov test, *P* = 0.57 for ASD; AUC = 0.57, *P* = 0.83 for ADHD; AUC = 0.55, *P* = 0.15 for MDD) (Fig. 3b-d). These results suggest that on the biological dimension defined by the SSD classifier, ASD, ADHD and MDD were not close to SSD.

**Fig. 3.**
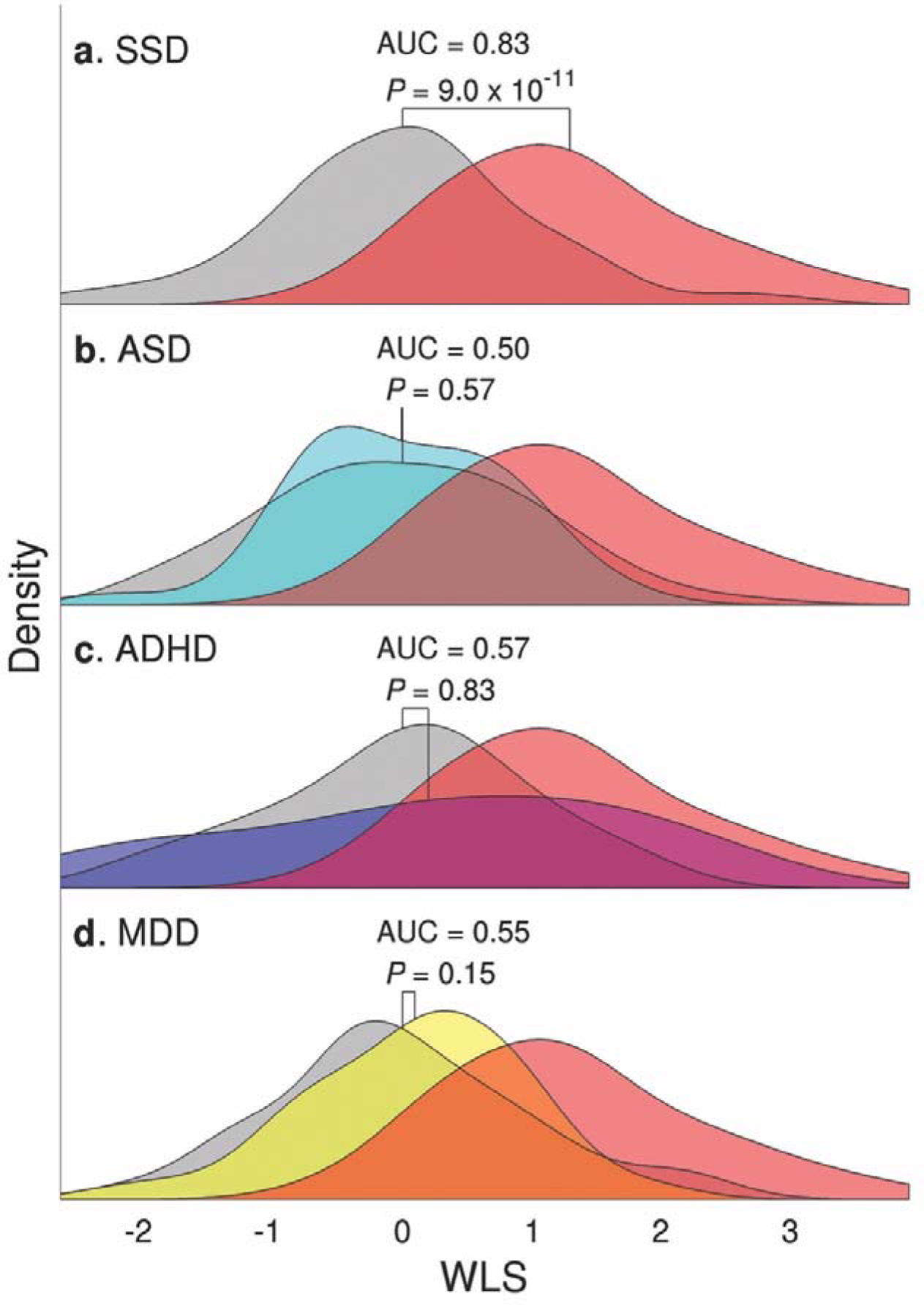
Application of the SSD classifier to other psychiatric disorders (ASD, ADHD, MDD). The density distribution of the weighted linear summation (WLS) was obtained by applying the SSD classifier to (**a**) SSD, (**b**) ASD, (**c**) ADHD, (**d**) MDD datasets. In each panel, patient distribution and HC distribution are plotted separately, with colored (SSD: red; ASD: cyan; ADHD: blue; MDD: yellow) and grey (HC) areas, respectively. For reference, WLS distribution of the SSD patients in **A** is duplicated across the panels (**b-d**). For each patient and control pair in (**a-d**), statistical significance was tested by Benjamini-Hochberg-corrected Kolmogorov-Smirnov.

### Relationships between SSD and ASD on the two biological dimensions

There were two main findings of relationships between SSD and ASD (Fig. 4a and Supplementary Results). First, the center of the SSD population on ASD dimension was elevated to near 0.5 with respect to the center of its HC population, while the center of the ASD population on SSD dimension remained at zero, the same as the center of its HC population. Second, the SSD and ASD dimensional scores were significantly correlated in the ASD population (*r* = 0.28, *P* = 0.040, permutation test corrected for multiple comparisons), while there was no correlation in the SSD population. Most of the ASD classifier’s FCs consistently contributed to the SSD-HC classification, but the FCs selected by the SSD classifier made inconsistent contributions to the ASD-HC classification, resulting in a cumulative WLS close to zero (Fig. 4b and Supplementary Results). The first asymmetry finding was interpreted by the differences of contribution results. The cumulative sum of the correlation coefficients within the ASD population indicated a general positive trend (Fig. 4c and Supplementary Results), which was the same direction as the largest correlation. On the other hand, the sum of the correlation coefficients within the SSD population was close to zero, due to contradicting correlation coefficients. The second asymmetry finding was explained by these correlation coefficients analysis. Moreover, the number of FCs selected across LOOCV of the HC-SSD classification was more than double that of HC-ASD, and the FC with the largest absolute weight in the SSD classifier (FC_1_^SSD^) was selected in only 15% of the total LOOCV folds. These results were summarized in the schema (Fig. 4d and Supplementary Results).

**Fig. 4.**
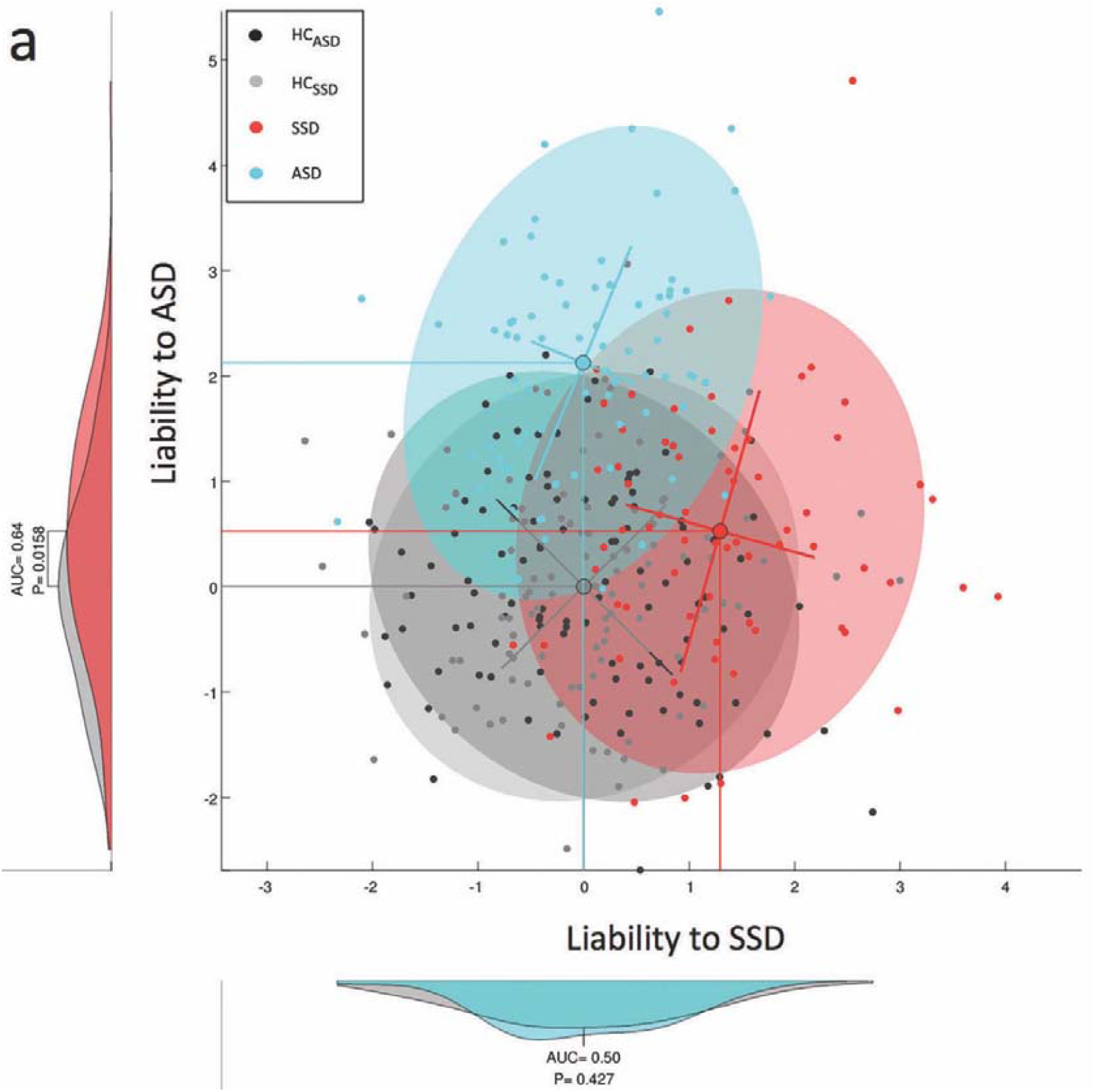
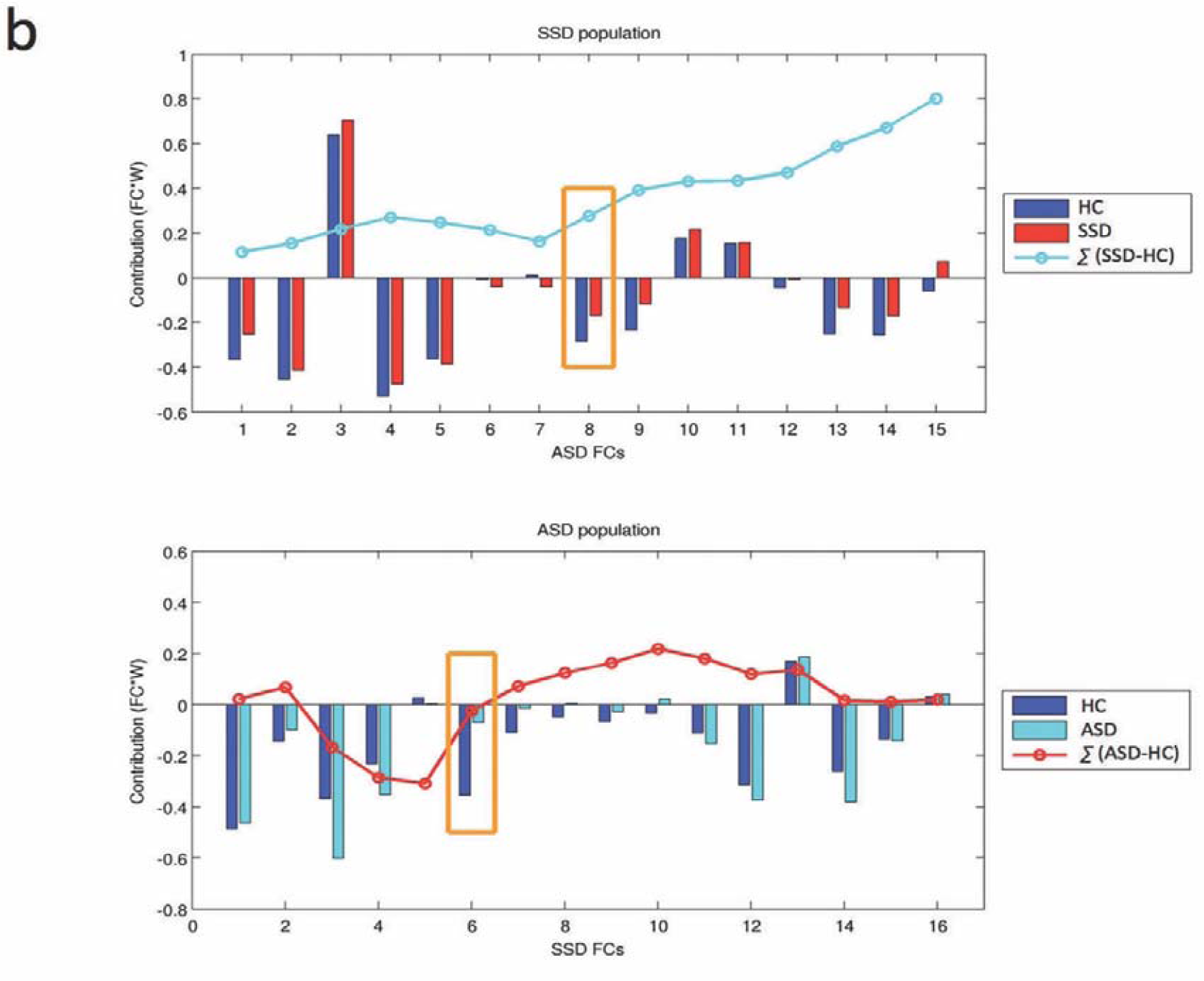
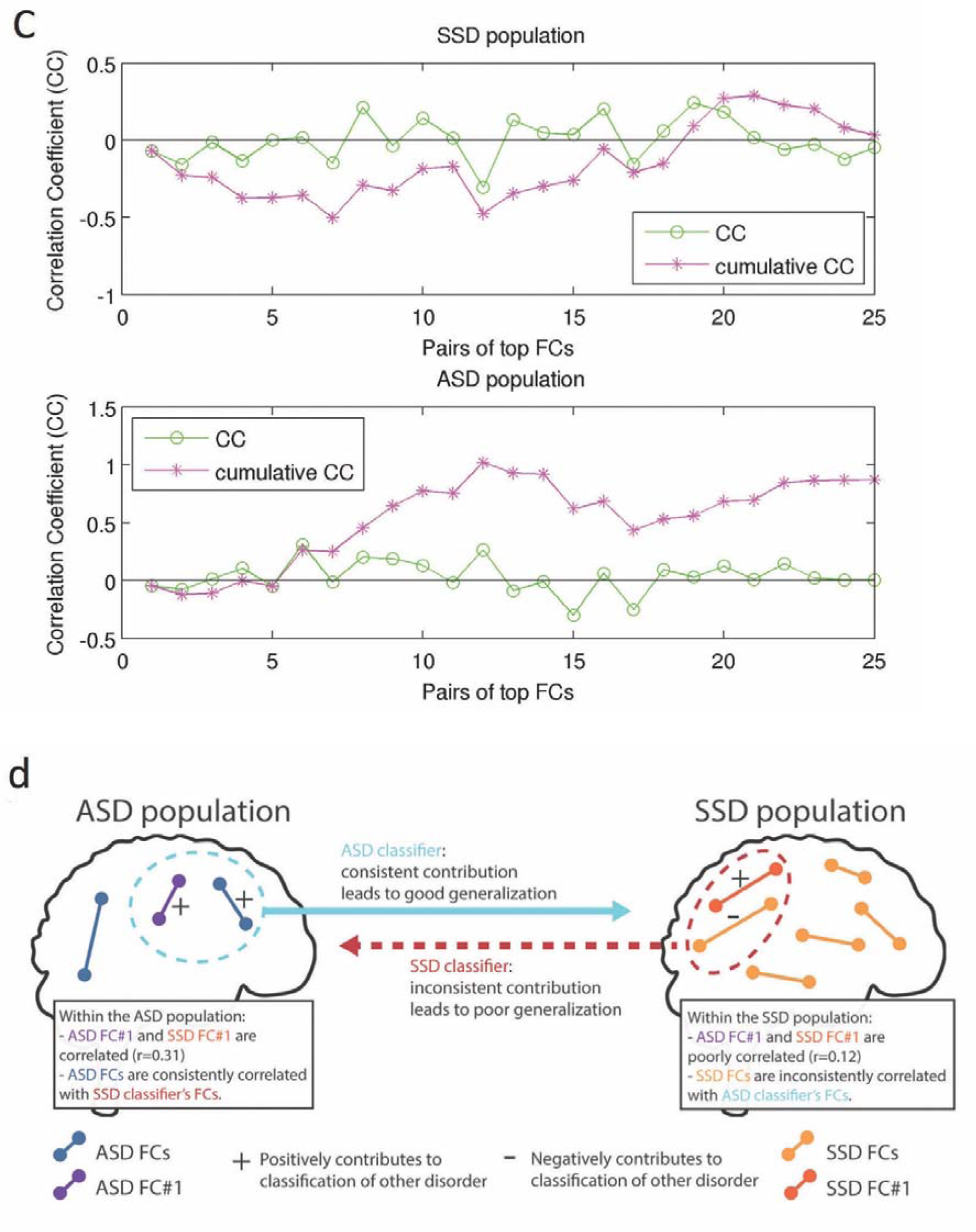
Relationships between SSD and ASD on the two biological dimensions. (**a**) Individuals with SSD, ASD, and HC on the SSD-ASD dimension plane. On the abscissa, the SSD dimension is the weighted linear summation (WLS) computed using the SSD classifier. On the ordinate, the ASD dimension is the WLS computed using the ASD classifier. The WLS of each dataset was normalized so that controls have zero mean and unit variance (statistical analysis was not affected by this normalization). (**b**) Contribution of each FC of a classifier to the WLS of the alternative disorder. The upper and lower graphs are the contributions of the ASD classifier’s FCs to the SSD-HC discrimination and the SSD classifier’s FCs to the ASD-HC discrimination, respectively. A large positive difference between SSD (red column) or ASD (cyan column) and HC (blue column) contributions indicates that a specific FC contributes positively to the classification. The cumulative sum (cyan or red line) of all the differences shows the contribution of each FC to the total WLS. The orange-highlighted box showed the FC with the largest absolute weight in the ASD classifier (FC_1_^ASD^) and in the ASD classifier (FC_1_^SSD^). (**c**) Correlation coefficients between the top 5 FCs of the SSD and ASD classifiers. The correlation coefficients (CC) between each pair of top FCs were computed (green line). The cumulative correlation coefficient (pink line) is computed in order to analyze the general trend of correlation, and a larger value indicates that the majority of the pairs have a positive correlation. The upper and lower graphs show pairs of top FCs within the SSD and ASD populations, respectively. (**d**) Schema of the relationship between ASD and SSD. The represented location of the FCs does not correspond to their true location. The FCs within the dashed circles represent those selected also by the final one-shot classifier, while those outside are those selected during LOOCV. FC_1_^ASD^=ASD FC#1 is the FC with the largest absolute weight in the ASD classifier (FC_1_ASD: right thalamus – left subcallosal sulcus). FC_1_^SSD^=SSD FC#1 is the FC with the largest absolute weight in the SSD classifier (FC_1_SSD: left central sulcus – right calloso marginal anterior fissure).

## Discussion

To our knowledge, this is the first study to quantify overlapping, but asymmetrical relationships between SSD and ASD, by combining the categorical approach based on DSM and the dimensional approach based on brain connectivity. The sophisticated machine-learning algorithms using categorical diagnostic labels and whole brain rs-fMRI produced a classifier that could discriminate patients from HC. At the same time, the classifier generated a probabilistic degree of liabilities to SSD and ASD based on whole brain functional connectivity from the WLS distributions. The neural liability was so continuous that we could regard it as a biological dimension. Moreover, the biological dimension needs to be robust enough to have generalizability to independent cohorts, as the biological dimension should be compatible with diagnoses that are common in different cohorts. Here, we developed the SSD classifier by a similar method to that described for our previous ASD classifier ^30^. The SSD classifier had generalizability to two independent cohorts in different countries and MRI machine vendors, not to other psychiatric disorders, and had specificity to chronic patients. Using these two classifiers, we could visualize individuals with ASD and SSD with their relative liability, and determine the overlapping, but asymmetrical relationships between SSD and ASD populations on the two biological dimensions. The relationships were more complicated than previously discussed in conceptual frameworks ^2^.

The ASD classifier was developed in our previous study ^30^, and here we focused on generating the SSD classifier. Various machine-learning algorithms have been applied previously to develop SSD classifiers that could discriminate patients with SSD from HC ^34-39^. However, none of the previous studies using only rs-fMRI tested whether the classifiers could present generalizability across different countries and MRI machine vendors. It was reported that there was a significant effect of MRI machine vendors ^40^ and ethnicities on MRI signals ^41^. A robust universal classifier should have generalizability to cohorts in a range of different countries under varying scanning protocols and imaging apparatus. Our classifier achieved high AUC (generalizability) to COBRE and UMCU-TOPFIT over the differences of various countries and MRI machine vendors. In contrast to COBRE and UMCU-TOPFIT, the SSD classifier achieved lower AUC (0.42) for a JHU-FES dataset. This can be attributed to the differences in the patients’ disease stage. Indeed, previous studies reported consistent differences in FC patterns between chronic SSD and FES ^42, 43^. Consequently, the finding that the SSD classifier did not generalize to FES might indicate that the classifier was specific to patients at a chronic stage of disease. In addition, we confirmed the specificity of the SSD classifier by demonstrating that it did not discriminate other psychiatric disorders from their respective control populations.

Plotting individuals with ASD, SSD, and HC on the dimensions along with DSM could show their heterogeneity based on functional neural circuits. A dimensional approach from only biological features using machine-learning algorithms could identify biotypes, but the biotypes were far from clinical diagnoses ^13^. In contrast, our biological dimensions were compatible with DSM, and the continuous WLS distributions of ASD and SSD populations were matched to the current psychiatric approach in DSM known as “spectrum”. Thus, our combined method of biological dimensions and the DSM system in this study may be useful in daily clinical work. Identifying a patient on the SSD-ASD dimensions may contribute to a clinician’s medical decision-making.

Several alternate models about the relationship between ASD and SSD have been proposed ^2^. While these models were within conceptual frameworks, some studies that applied biological methods actually showed commonalities ^4, 7^, or diametric conditions ^44, 45^ between the two disorders. We took advantage of the two biological dimensions of ASD and SSD, and revealed an overlapping, but asymmetrical relationships, which cannot be attained by a single dimension. The asymmetries here have dual meanings. First, the SSD population showed increased liability on the ASD dimension, while the ASD population did not on the SSD dimension. Increased ASD liability in the SSD population contributed to the substantial overlap between SSD and ASD populations (Fig. 4a). Second, the two dimensions were correlated within the ASD population but not in the SSD population. The results from LOOCV underlying these asymmetries suggested that the network SSD is characterized by a larger diversity and that it partially shares information with the smaller network of ASD. This is in agreement with recent genetic evidences that ASD shares a significant degree of polygenic risk with SSD ^4^, and that common genetic variations explain nearly 50% of total liability to ASD ^46^ and 25-33% of total liability to SSD ^47^, suggesting that environmental factors play a significant role in the heterogeneous etiopathogenesis of schizophrenia ^48^.

### Limitations

First, the AUC of UMCU-TOPFIT (0.66) was lower than the AUC of COBRE (0.75). There was a difference in MRI raw data between 3D scan in UMCU-TOPFIT and 2D scan in COBRE. The classifier was developed from Kyoto datasets in 2D scan, and this might be related to the AUC difference. Second, almost all patients were on antipsychotic medication. Previous studies reported that antipsychotics altered the functional connectivity in frontal and striatal circuits ^49, 50^. Although we found no significant correlation between the SSD classifier and antipsychotic medication (Supplementary Results), potential effects of antipsychotics on the SSD classifier cannot be entirely ruled out. Third, we did not recruit comorbid patients (ASD with psychosis), and we did not discuss comorbidity.

### Conclusion

The current findings obtained by the two biological dimensions consisting of functional connectivity revealed asymmetrical and overlapping relationships between SSD and ASD.

## Acknowledgements

We thank the patients and controls for participating in these studies. This research was conducted as the “Application of DecNef for development of diagnostic and cure system for mental disorders and construction of clinical application bases” of the Strategic Research Program for Brain Sciences from the Japan Agency for Medical Research and Development, AMED. The UMCU-TOPFIT study received funding from the Dutch Diabetes Research Foundation (2007.00.040), Lilly Pharmaceuticals, Houten, The Netherlands (Ho01-TOPFIT); Janssen Pharmaceuticals, Tilburg, The Netherlands; and the Dutch Psychomotor Therapy Foundation, Utrecht, The Netherlands. The JHU FES study was supported by the National Institute of Mental Health (NIMH) grant number MH-094268, MH-092443, MH-105660, and the Silvio O. Conte center funded by NIMH, the National Institutes of Health grant number P41EB015909 and R01NS084957, grants from Stanley, S-R/RUSK, and NARSAD, and parts of the participant recruitment was supported by Mitsubishi Tanabe Pharm. Co. Ltd. (USA).

